# Whole-genome sequencing identifies interferon induced protein IFI6 as a strong candidate gene for VNN resistance in European sea bass

**DOI:** 10.1101/2022.05.31.494209

**Authors:** Emilie Delpuech, Marc Vandeputte, Romain Morvezen, Anastasia Bestin, Mathieu Besson, Joseph Brunier, Aline Bajek, Boudjema Imarazene, Yoannah François, Olivier Bouchez, Xavier Cousin, Charles Poncet, Thierry Morin, Jean-Sébastien Bruant, Béatrice Chatain, Pierrick Haffray, Florence Phocas, François Allal

**Affiliations:** MARBEC, Univ. Montpellier, Ifremer, CNRS, IRD, INRAE, 34250 Palavas-les-Flots, France; Université Paris-Saclay, INRAE, AgroParisTech, GABI, 78350 Jouy-en-Josas, France; SYSAAF, Station LPGP/INRAE, Campus de Beaulieu, 35042 Rennes, France; Ecloserie Marine de Gravelines-Ichtus, Gloria Maris Groupe, 59273 Gravelines, France; Fermes Marines Du Soleil, 17840 La Brée Les Bains, France; ANSES, Viral Fish Diseases Unit, Technopôle Brest- Iroise, 29280 Plouzané, France; US 1426, GeT-PlaGe, INRAE, Genotoul, Castanet-Tolosan, France; INRAE-UCA, UMR GDEC, 63000 Clermont-Ferrand, France

## Abstract

**Background:** Viral Nervous Necrosis (VNN) is major disease affecting of European sea bass. Understanding the biological mechanisms that underlie VNN resistance is thus important for the welfare of farmed fish and the sustainability of production systems. This study aimed at identifying key genomic regions and genes that determine VNN resistance in sea bass.

**Results:** We generated a dataset of around 900,000 single nucleotide polymorphisms (SNPs) identified from whole-genome sequencing (WGS) in the parental generation in two different commercial populations (pop A and pop B) comprising 2371 and 3428 European sea bass with phenotypic records for binary survival in a VNN challenge. In each commercial population, three cohorts were submitted to the redspotted grouper nervous necrosis virus (RGNNV) challenge by immersion and genotyped on a 57K SNP chip. After imputation of WGS SNPs from their parents, QTL mapping was performed using a Bayesian Sparse Linear Mixed Model (BSLMM). We found several QTL regions on different linkage groups (LG), most of which are specific to a single population, but a QTL region on LG12 was shared by both commercial populations. This QTL region is only 127 kB wide, and we identified IFI6, an interferon induced protein at only 1.9 kB of the most significant SNP. An unrelated validation population with 4 large families was used to validate the effect of the QTL, for which the survival of the susceptible genotype ranges from 39.8 to 45.4%, while that of the resistant genotype ranges from 63.8 to 70.8%.

**Conclusions:** We could precisely locate the genomic region implied in the main resistance QTL at less than 1.9 kb of the interferon alpha inducible protein 6 (IFI6), which has already been identified as a key player for other viral infections such as hepatitis B and C. This will lead to major improvements for sea bass breeding programs, allowing for greater genetic gain by using marker-assisted genomic selection to obtain more resistant fish. Further functional analyses are needed to evaluate the impact of the variant on the expression of this gene.

## Background

The sustainable development of aquaculture is strongly dependent on the capacity to control the epidemics that can impact the farms [1]. Viral Nervous Necrosis (VNN), also called viral encephalopathy and retinopathy (VER), is caused by Nervous Necrosis Virus (NNV), a single-stranded positive-sense RNA virus. It is a major threat to the marine and freshwater fish farming industry worldwide causing increased mortality, degradation of feed conversion ratio and of animal welfare, thus affecting both production and economic viability of farms [2]. Among the many species affected by this disease [3], the European sea bass can be highly impacted in the Mediterranean aquaculture industry, due to a warm environment compatible with the occurrence of VNN [3–6]. This disease can devastate an entire production of particularly susceptible larvae and juveniles, with a mortality rate that can reach 90% or more, while adults can also be impacted [3, 7]. Infected fish are mainly recognizable by their abnormal swimming behaviour [8]. They show severe neurological disorders due to intensive vacuolation of the retina and the nervous system [9], and a fast mortality peak, usually on the 10th day after infection with the NNV [10].

Vaccination studies were conducted for this disease, and some recently obtained very positive results [11] but cannot yet be applied on larvae or early juveniles, for which vaccination remains a challenge [12, 13]. Genetic improvement remains an important potential way to improve disease resistance in aquaculture in general [14] and specifically to improve VNN resistance [15]. Recent studies have shown that there is genetic variation in resistance to VNN in European sea bass with a heritability ranged between 0.26 and 0.43 [10, 16–19]. After validation of the possibility to select and improve fish for resistance to NNV [14], dissecting the genetic architecture of VNN resistance in several European sea bass populations is a necessary step to obtain a better understanding of the molecular architecture of infection-resistant fish and to permit potential marker-assisted selection [20]. For this, genome-wide association studies (GWAS) have become a powerful tool in genetics and commonly applied in animal breeding [21, 22] e.g. in different sea bass species and populations for VNN resistance [10, 18, 19, 23]. They reported the identification of quantitative trait loci (QTL) and even the identification of SNP variants impacting the survival rate of sea bass exposed to NNV. In particular, a QTL was identified on the LG12 in several populations of European sea bass, indicating a genomic region of interest for improving VNN survival [10, 18]. However, from these studies the localisation of this QTL remained unprecise and population dependent, while the implementation of efficient marker assisted selection requires identifying the causal variant(s) or very tightly linked loci. With this aim, we imputed whole genome sequence on 5779 sea bass from two French breeding programs challenged for VNN, and performed a GWAS with a Bayesian Sparse Linear Mixed Model (BSLMM) to identify the main genomic region of interest, which was then validated in an independent population.

## Methods

### Ethic statement

All infection challenges were carried out in accordance with the European guidelines (Directive 2010–63-EU) and the corresponding French legislation. Animal experiment procedures were approved by the ethics committee on animal experimentation COMETH ANSES/ENVA/UPC No.16 and were authorized by the French Ministry of Higher Education, Research and Innovation under numbers 2017022816255366, 29/01/13-5 and 10/03/15-1.

### Populations and experimental design

#### Discovery populations

A total of 5799 European sea bass (*Dicentrarchus labrax*) from commercial populations were used in our study to identify the QTLs. These animals produced by artificial mating, come from two different French hatcheries designated as pop A and pop B. The 2371 individuals from pop A were distributed in three cohorts and the 3428 individuals from pop B were also distributed in 3 cohorts. So, six cohorts were used in this study, including two cohorts that were previously studied [10]. To identify the cohorts in the commercial populations, an additional number from 1 to 3 will be added to pop A or pop B.

In pop A, the three cohorts were not related and these individuals were distributed as follows: 671 individuals from cohort A_1, deriving from a partly factorial mating of 56 sires and 19 dams, 650 individuals from cohort A_2, deriving from a partly factorial mating of 58 sires and 16 dams and 1050 individuals from cohort A_3 deriving from a partly factorial mating of 60 sires and 20 dams. All 174 sires of these individuals were sampled for genome sequencing. In pop B, cohorts 1 and 2 were related while cohort 3 was more distantly related. Pop B cohorts included 1083 individuals from cohort B_1, deriving from a partly factorial mating of 40 sires and 14 dams, 1087 individuals from cohort B_2, deriving from a partly factorial mating of 41 sires and 15 dams and 1258 individuals for cohort B_3 deriving from a partly factorial mating of 48 sires and 27 dams. For this population, parents of cohorts B_2 and B_3 were all sampled for sequencing and for cohort B_1 some pairs of parents were sampled but for a portion of individuals only one parent (mostly the mothers) had enough DNA to be sequenced. In the end, 159 parents from pop B were sampled for whole-genome sequencing. All this information is reported in table 1.

**Table 1:**
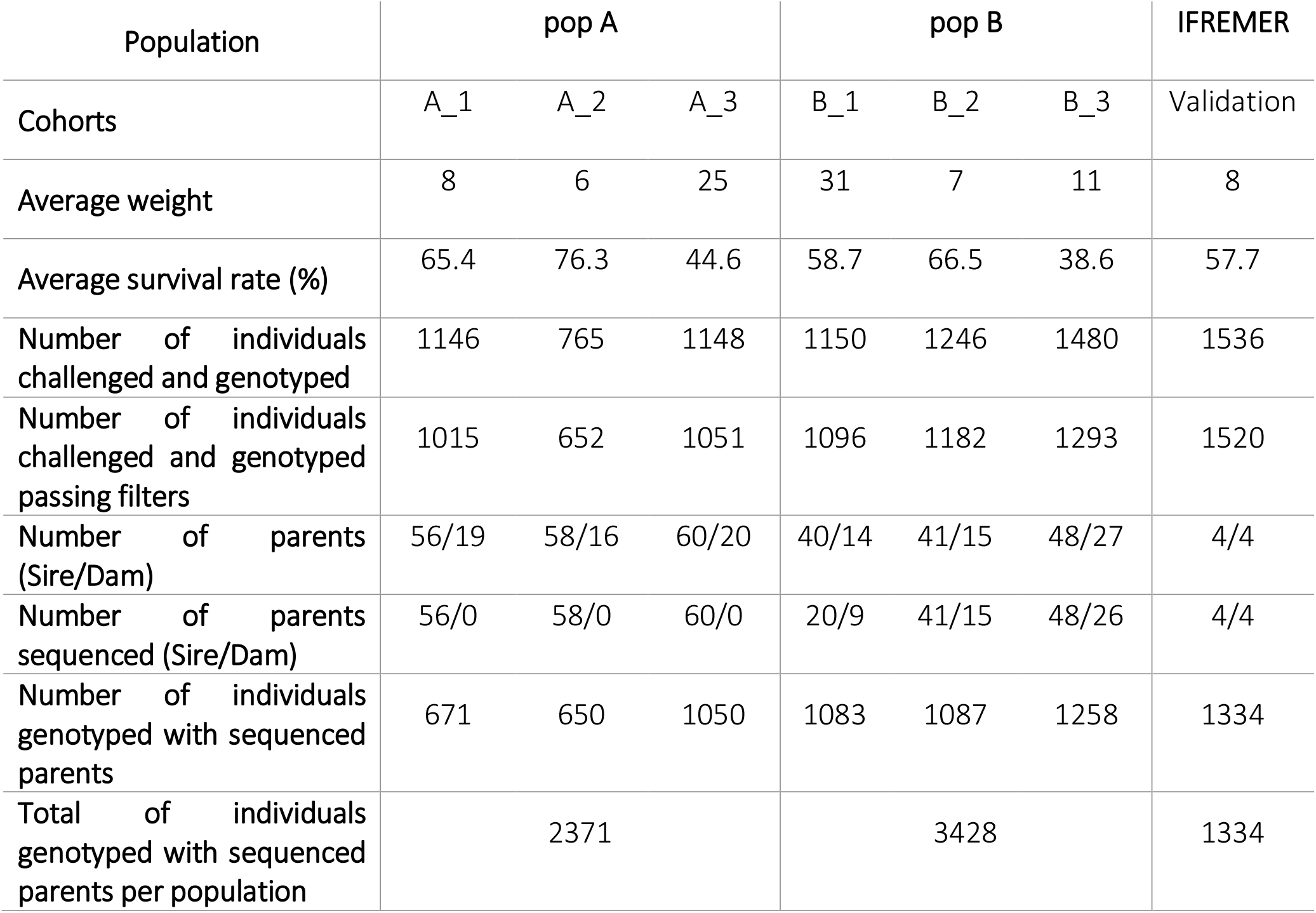
Description of the sampling strategy applied for each of the two commercial populations and for the validation population challenged for VNN.

European sea bass from the commercial cohorts were challenged for VNN on the SYSAAF-ANSES Fortior Genetics platform (ANSES, Plouzané, France). For all challenges, fish were maintained in filtered seawater at a temperature of 27 °C ± 2 °C in a flow-through system. Infectious challenges were performed separately but in the same manner for all cohorts. All fish sent to the infectious challenge were individually tagged with RFID glass tags. The fish were acclimatized for a period of three weeks before the challenge was performed. To summarize, infection with the redspotted grouper nervous necrosis virus (RGNNV) strain W80 was performed in a static bath of aerated seawater containing 1 × 10^5^ TCID_50_/ml of the virus.

Mortality was recorded daily during the challenge period which was 27 days for pop A and 42 days for pop B. To confirm the presence of the NNV virus in dead fish and eliminate the possible impact of unwanted bacterial coinfections, bacteriological and viral analyses were performed during the challenges and during the peak of mortality.

#### Validation population

For the validation part, four experimental full-sib families challenged for VNN were used. These families have already been described and studied by Griot et al. 2021 [10], namely NEM10, NEM12, SEM8 and WEM18. These individuals were then challenged for VNN following the same method used for the commercial populations. In total, 1536 European sea bass from these families were challenged for NNV resistance for 33 days. Its challenge results were used for the validation of the results obtained from the commercial cohorts. So, statistically these four experimental full-sib families constitute the validation population. All eight parents of these four families were sampled for whole genome sequencing.

### Whole-genome sequencing and variant calling

A total of 333 parents of individuals challenged for resistance to VNN were collected to obtain whole-genome sequencing (WGS) data. For commercial populations, 174 parents from pop A and 159 parents from pop B were sequenced. In addition, the 8 parents of the validation population were also sequenced (see details in Table 1). The genomic DNA was extracted using the standard phenol–chloroform protocol from the parents’ pectoral fin biopsies at the GeT-PlaGe core facility, INRAE Toulouse, to perform DNA sequencing. DNA-seq libraries were prepared according to Illumina’s protocols using the Illumina TruSeq Nano DNA HT Library Prep Kit. Briefly, DNA was fragmented by sonication, size selection was performed using SPB beads (kit beads), and adaptors were ligated for traceability and sequencing. Library quality was assessed using a Advanced Analytical Fragment Analyzer and libraries were quantified by QPCR using the Kapa Library Quantification Kit. DNA-seq experiments were performed on an Illumina NovaSeq6000 using a paired-end read length of 2×150 pb with the Illumina NovaSeq6000 Reagent Kits.

The obtained sequencing reads were aligned to the European sea bass reference genome consisting of 24 linkage groups (LG) (seabass_V1.0) using the Burrows-Wheeler Aligner (BWA, v.2.1) method with default parameters [24]. Duplicates were marked with Picard (http://broadinstitute.github.io/picardv.2.21.1). After mapping, SNPs and InDels were called using the DeepVariant (v.1.1.0) tool [25] to keep all types of variants present in the analyzed populations. Next, the dataset identifying SNPs with quality greater than 30 and with a coverage higher than 4 was kept using BCFtools (v.1.13) [26]. A second step of filtering was performed with the PLINK software (v.1.9) [27, 28] to remove SNPs with: (1) a missing call rate higher than 10%, (2) a minor allele frequency (MAF) lower than 1% and (3) more than two alleles identified. Finally, the output VCF files were converted to plink format and were coded as 0, 1 and 2 corresponding respectively, to the homozygous genotype for the reference allele from the published genome (seabass_V1.0), heterozygous genotype and homozygous genotype for the alternative allele.

### SNP chip genotyping and imputation to whole-genome sequences

Samples from challenged cohorts were genotyped for 56,730 SNPs using the ThermoFisher Axiom™ Sea Bass 57k SNP array DlabChip, at the Gentyane genotyping platform (INRAE, Clermont-Ferrand, France). A total of 3059 and 3876 individuals were genotyped in pop A and pop B, respectively. In the validation population, a total of 1536 individuals were also genotyped with the same chip. Then, SNP calling was done using ThermoFisher’s AxiomAnalysisSuite™ software to identify genotypes at each probe set. Preliminary quality controls were applied with cutoff values of 95% for SNP calling rate,90% for sample calling, and the “Run PS Supplemental” option was used to regenerate SNP Metrics to select SNPs identified polymorphic by the software. With the PLINK software, two other filtering were applied on genotypes of challenged individuals within commercial populations: remove SNPs with a MAF less than 5% and SNPs with a p-value for the Hardy-Weinberg test less than a threshold of 10^−8^. The output VCF files were converted to plink format and were coded as 0, 1 and 2 corresponding respectively, to reference allele from the published genome (seabass_V1.0), heterozygous and alternative allele. Parentage assignment was performed using 1000 sampled markers (with a MAF around 0.5) with the APIS R package [29] with a positive assignment error rate set at 1%, to assign all challenged sea bass to their sequenced parents.

Imputation of the WGS SNPs to the offspring genotyped with the 57k SNP array DlabChip was performed using the FImpute software (v.3) [30], accounting for pedigree information. SNPs identified from the WGS variant calling of the parents were used as reference and the 57K genotypes of the challenged offspring were used as target in each population analyzed (pop A, pop B and validation population).

To perform the most efficient approach in both commercial populations, a subset of the imputed WGS SNPs was created. Two successive filters were applied to be on the same set of SNPs across populations and to avoid redundancy among SNPs. First, we retained common SNPs between the two commercial and the validation populations, secondly, we filtered within population the remaining SNPs based on linkage disequilibrium calculated between all pairs of SNPs with an upper limit set to r^2^=0.8 within a 200 kb windows in PLINK; and finally reselecting common SNPs after filtering on linkage disequilibrium in each population.

### Bayesian sparse linear mixed model for genome wide association studies

GWAS was based on a Bayesian Sparse Linear Mixed Model (BSLMM) that assumes that all SNPs have at least a relatively small effect, but also that a few SNP may have a large effect [31]. Therefore, BSLMM is capable of adapting to different genetic architectures of the studied trait, from the infinitesimal polygenic model to a model which assumes that only a very small proportion of all variants affect the phenotype. The general model can be presented as follows:

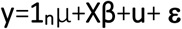

where 1_n_ is an n-vector of 1s, µ is a scalar representing the phenotype mean, **X** is an n × p matrix of genotypes measured on n individuals at p SNPs, **β** is the corresponding p-vector of the SNP effects; **u** is a vector of random additive genetic effects distributed according to ***N***(0, **K**σ^2^_b_), with σ^2^_b_ the additive genetic variance and **K** the genomic relationship matrix; and **ε** is a n-vector of residuals ***N***(0, **I** σ^2^_e_), σ^2^_e_ is the variance of the residual errors. Assuming **K** = **XX**^T^ /p, the SNP effect sizes can be decomposed into two parts: α that captures the small effects that all SNPs have, and β that captures the additional effects of some large effect SNPs. In this case, u = Xα can be viewed as the combined effect of all small effects, and the total effect size for a given SNP is **γ**_i_ =α_i_ + β_i_. The individual SNP effects **γ**_i_ are sampled from a mixture of two normal distributions, **γ**_i_∼πN(0,σ^2^_a_+σ^2^_b_)+(1−π)N(0,σ^2^_b_) where σ^2^_b_ is the variance of small additive genetic effects, σ^2^_a_ is the additional variance associated to large effects and π is the proportion of SNPs with large effects.

In our study, the BSLMM was implemented using Genome-Wide Efficient Mixed Model Association (GEMMA) software [31]. For a more accurate study of the VNN survival trait, this binary trait was adjusted by the cohort effect in each population effect using a linear model.

Within each population, the GWAS was performed by combining data from the 3 cohorts to increase the power of the analysis. In GEMMA, we applied a linear BSLMM model using MCMC as proposed by Zhou et al [31] for the survival trait. A total of 5 million iterations (-s option) were performed with a burn-in of 100,000 steps and we recorded one state in every 10 iterations for further analysis. In addition, to ensure convergence of the distribution of the hyper-parameter π, the minimum and maximum numbers of SNPs that was sampled to be included be included in the model were set to 5 (-smin option) and 100 (-smax option), respectively. These threshold values were based on an initial run with default values indicating median numbers of about 20 SNPs as showing a large effect.

### QTL region and annotation

BSLMM uses a Markov chain Monte Carlo algorithm to sample from the posterior distribution to obtain all parameters values, among which the SNP effect estimates 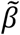, but also the hyper-parameter π and a posterior inclusion probability (PIP) for each SNP that indicates the proportion of samples in which that SNP is classified as having a large effect. This proportion can be used for QTL mapping as it indicates the strength of the evidence that the SNP has to be included in the model. In other words, SNPs that are most robustly associated with the phenotype are therefore expected to have large PIPs. Following Barbieri and Berger [32], regions with a PIP above 0.5 were selected (since SNPs from these regions are included in the model in the majority of iterations). As a complementary approach, to define a less stringent threshold, Stephens and Balding [33] proposed to calculate a Bayes Factor (BF) as follow :

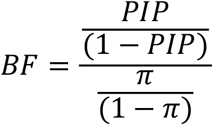

As proposed by Kass and Raftery [34], the natural logarithm transformation of the BF (logBF) was computed as twice the natural logarithm of the BF. This logarithmic scale produced values within the same range as usual likelihood ratio test values, thus facilitating the determination of thresholds to define QTLs. A threshold logBF ≥ 10 was used for defining very strong evidence for a QTL according to Michenet et al. [35]. The BSLMM results were visualized via a Manhattan plot where all negative values for logBF were set to 0. Credibility intervals were determined using the threshold logBF ≥ 10 for defining a peak SNP showing strong evidence for a QTL. The credibility interval included every SNP with a logBF > 5 within a 100 kb sliding window from the peak SNP. This procedure was repeated for each of the identified QTL regions.

To determine how QTL regions affected VNN resistance, the effects of SNPs characterizing each region on population survival rate were analyzed. For this purpose, we corrected the survival rate results by cohort in each population in order to be able to compare all challenges between them. Thus, we standardized all challenges to a survival rate of 50% after a probit transformation, and then, the percent survival of each genotype at a QTL was recalculated in each cohort. Finally, we averaged these survival rates by cohorts to facilitate the discussion of the results.

The next step was to perform a functional analysis of the QTL region identified, to link variants detected in the whole-genome sequencing of parents with the annotation of the European sea bass. All annotated genes positioned in the QTL region were reported. To understand the role of the variants in the coding regions of identified genes, all SNPs identified from the variant calling were annotated based on the UCSC annotation [36] of the European sea bass reference genome with SnpEff software (v.5.0) and default parameters [37]. SnpEff annotates variants in the VCF files based on their position and the reference annotation. The polymorphisms were classified per variant type as coding or non-coding, then the impact of each was identified from high to low, and the functional class was assigned to them as non-synonymous or synonymous (silent) SNPs. As we know that the causative mutation may not be the most significant variant in GWAS analyses, for the non-coding regions, Ensembl Variant Effect Predictor (VEP) software of Ensembl was used to understand the role of variants positioned between genes [38]. The VEP provides methods for a systematic approach to annotate and prioritize variants in large-scale sequencing analysis, using transcripts, regulatory regions, frequencies from previously observed variants, citations, clinical significance information, and predictions of biophysical consequences of variants.

## Results

### VNN phenotypes

The six challenges were conducted up to 40 days for the six commercial cohorts. NNV presence was confirmed by viral analyses on several subsets of dead fish during the infection, on which the absence of significant bacterial coinfection was also confirmed. Survival rates ranged from 44.5% to 76.3% for pop A cohorts and from 38.6% to 66.5% for pop B cohorts. For the validation population, the average survival rate was 56% (Table1). The peak of mortality was between 5 and 15 days after infection for all seven commercial and validation cohorts (Figure 1).

**Figure 1:**
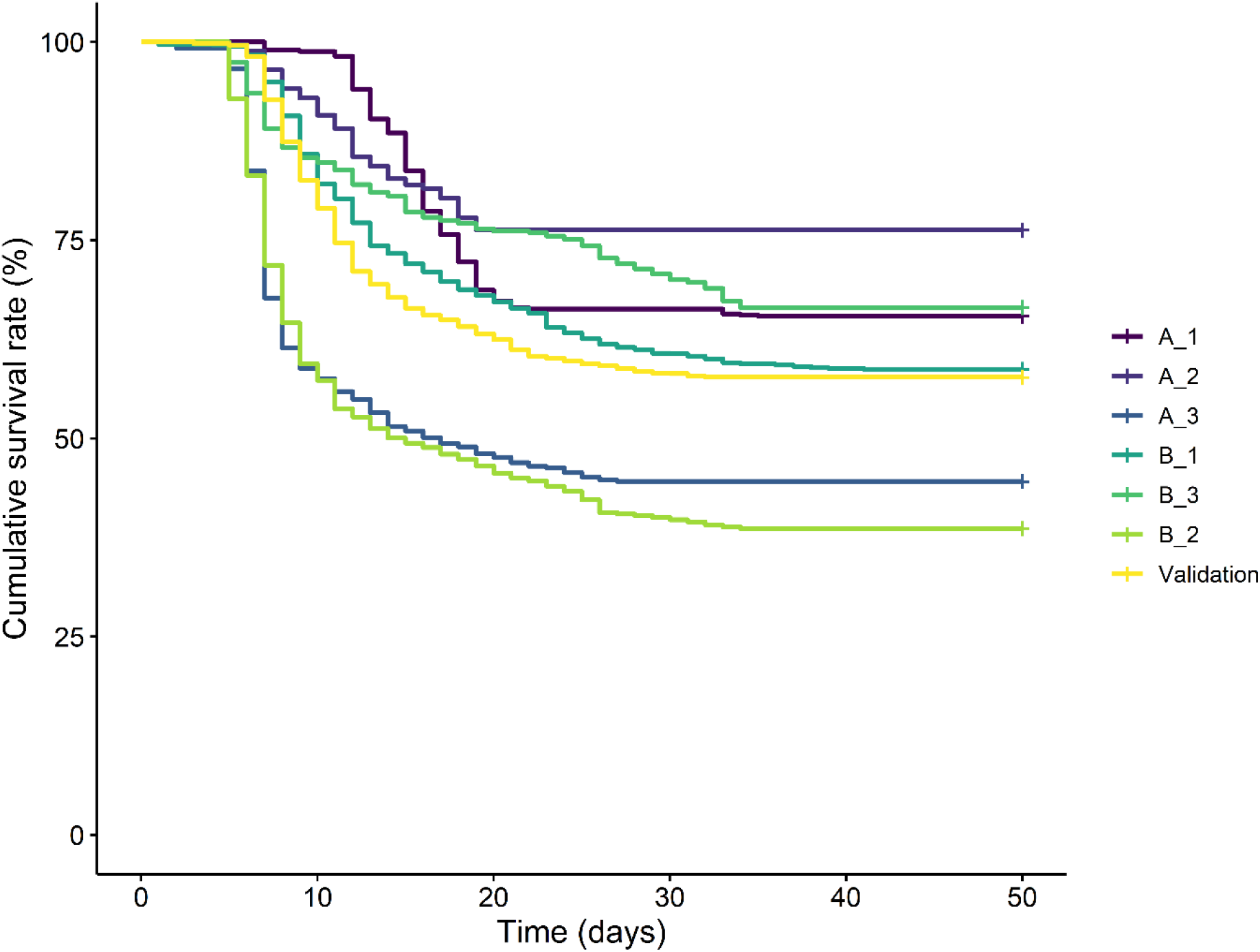
Evolution of the Kaplan-Meier cumulative survival rate for each of the 3 cohorts characterizing the two commercial populations, via a purple gradient for pop A (A_1, A_2, A_3) and a green gradient for pop B (B_1, B_2, B_3), during their respective VNN infection challenge. The results for the validation population are also plotted in yellow.

### From whole-genome sequences to imputed whole-genome variants

We generated WGS at an average coverage of 19-fold for 174 sires of pop A, 159 parents (126 sires and 53 dams) of pop B and 8 parents (4 sires and 4 dams) of the validation population. Variant calling data were available for all these fish and identified around 8 million raw variants. The quality control of the WGS SNPs was first carried out for each commercial population and the validation population. All individuals sequenced and considered in this study were kept. The number of SNPs removed by MAF (lower than 10%), SNP call rate (lower than 95%) and Hardy-Weindberg (p-value>10^−8^) filters are reported in Table S1. In total, 2,506,457 WGS filtered SNPs were kept in the 174 sequenced animals from pop A, 2,436,691 WGS SNP were kept in the 159 sequenced animals from pop B and 2,392,123 WGS SNP were kept in the validation population.

From the 5799 57K genotypes obtained in pop A and pop B respectively, 688 and 277 animals were removed with a call rate threshold of 90% and based on pedigree data i.e. keeping only those individuals that were challenged and had at least one parent sequenced. After SNPs filtering, following the MAF (lower than 10%), call frequency filters (lower than 95%) and the Hardy–Weinberg equilibrium filter, 40,743 SNPs and 36,862 SNPs were kept. Thus, for the 57k SNP array DlabChip, genotypes of 2371 pop A animals for 40,743 SNPs and genotypes of 3428 pop B animals for and 36,862 SNPs were retained for further analyses.

After the imputation step for the commercial populations, we obtained 2,506,457 and 2,436,691 imputed WGS SNPs for 2371 and 3428 challenged animals in pop A and pop B, respectively. Based on an r^2^>0.8 in 200 kb windows, 1,457,298 WGS SNPs and 1,457,105 WGS SNPs were retained for pop A and pop B respectively. Finally, to perform GWAS on commercial populations, a common subset of SNPs after applying the LD filters performed in each population, was created and was characterized by 838,451 imputed WGS SNPs.

### QTL detection for VNN resistance

The estimates of the genomic heritability (h^2^) of binary survival on the observed scale were 0.22 in pop A and 0.21 in pop B, combining the 3 cohorts in each commercial population. The estimation of the proportion of genetic variance explained by the polygenic effect represented 61% of the total genetic variance in pop A and 62% in pop B. In the BSLMM association analysis, 11 QTL regions in pop A and 12 QTL regions in pop B were identified as having large effects on the VNN resistance, meaning very strong evidence was found for these QTL (Table 2). In pop A, QTL were detected with strong evidence in LG1A, LG3, LG4, LG7, LG8, LG12, LG15, LG22-25 and LGx. In pop B, QTL detected with strong evidence were positioned in LG1B, LG7, LG8, LG9, LG12, LG13, LG14 and LG15 (Figure 2).

**Table 2:** List of QTL regions identified in the BSLMM analysis ranked according to their position on the genome (LG=Linkage Group ; PIP=Posterior Inclusion Probability ; POPULATION= population identification of the QTL region).

| QTL-ID | LG | MIN-POS (bp) | MAX-POS (bp) | PIP | logBF | Population |
| --- | --- | --- | --- | --- | --- | --- |
| LG1A_QTL | LG1A | 10,107,908 | 10,111,686 | 0.02 | 10.3 | pop A |
| LG1B_QTL | LG1B | 13,152,351 | 13,885,885 | 0.04 | 10.9 | pop B |
| LG3_QTL | LG3 | 10,137,843 | 10,137,843 | 0.03 | 10.6 | pop A |
| LG4_QTL | LG4 | 23,334,300 | 23,618,451 | 0.22 | 15.2 | pop A |
| LG7_QTL_1 | LG7 | 7,828,421 | 7,828,421 | 0.03 | 10.6 | pop B |
| LG7_QTL_2 | LG7 | 21,062,741 | 21,062,741 | 0.03 | 10.9 | pop A |
| LG8_QTL_1 | LG8 | 1,423,728 | 3,198,660 | 0.38 | 16.4 | pop A/pop B |
| LG8_QTL_2 | LG8 | 20,789,708 | 22,055,049 | 0.05 | 12.0 | pop A |
| LG9_QTL | LG9 | 4,532,879 | 6,335,544 | 0.18 | 14.4 | pop B |
| LG12_QTL_1 | LG12 | 4,717,307 | 7,246,406 | 0.20 | 14.6 | pop A/pop B |
| LG12_QTL_2 | LG12 | 8,671,217 | 8,797,942 | 0.59 | 18.5 | pop A/pop B |
| LG12_QTL_3 | LG12 | 9,118,505 | 9,821,502 | 0.36 | 16.2 | pop A/pop B |
| LG12_QTL_4 | LG12 | 12,167,000 | 19,133,722 | 0.30 | 16.0 | pop A/pop B |
| LG13_QTL | LG13 | 11,019,998 | 11,019,998 | 0.15 | 14.0 | pop B |
| LG14_QTL | LG14 | 26,274,292 | 26,276,196 | 0.07 | 12.1 | pop B |
| LG15_QTL | LG15 | 5,885,758 | 5,885,758 | 0.04 | 11.0 | pop B |
| LG15_QTL | LG15 | 24,678,361 | 24,678,361 | 0.07 | 12.3 | pop B |
| LG22-25_QTL | LG22-25 | 4,185,377 | 4,185,377 | 0.03 | 10.5 | pop A |

**Figure 2:**
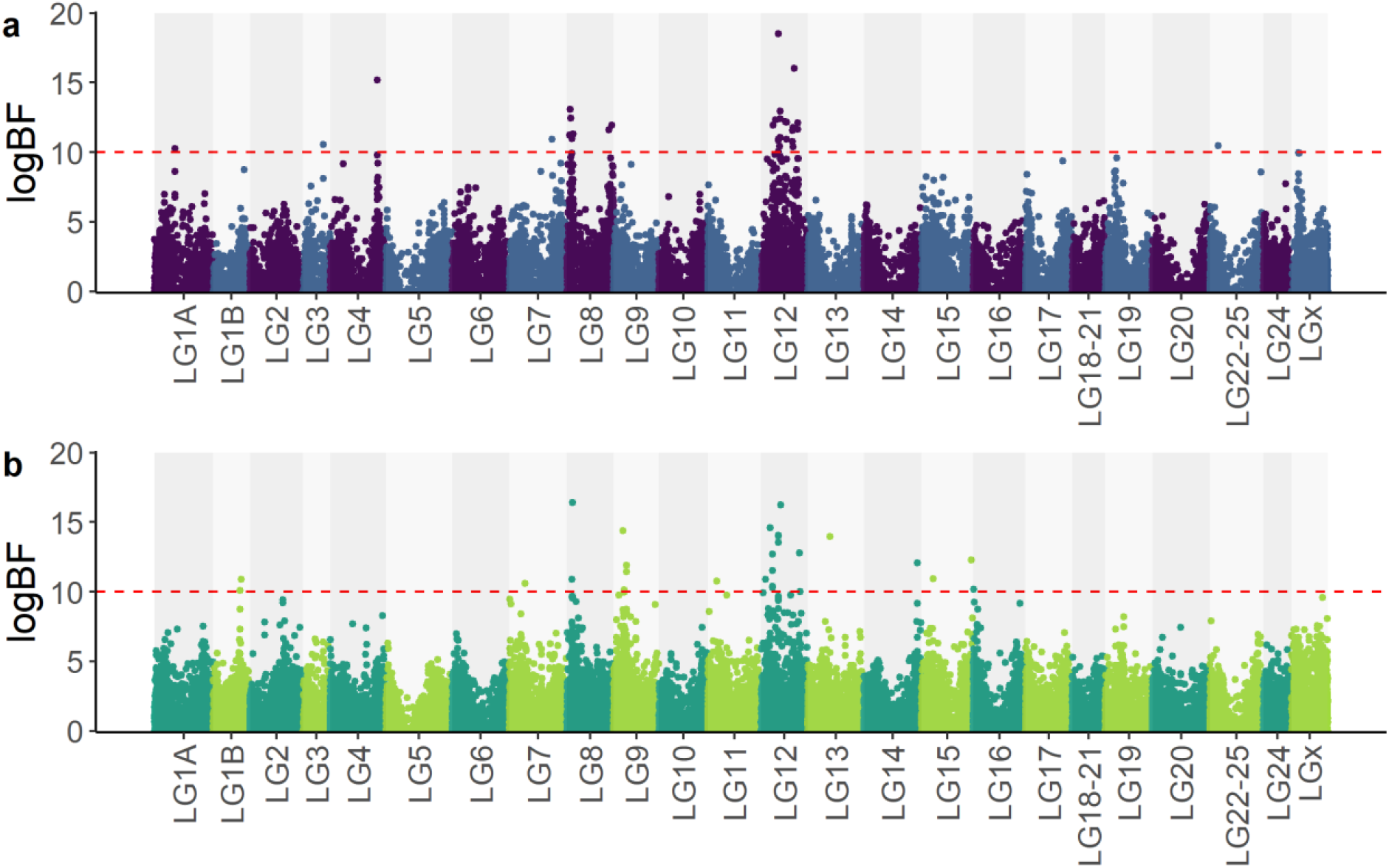
Manhattan plot of BSLMM results with logBF scale for VNN resistance in European sea bass populations pop A (a) and pop B (b). The red dashed line represents the logBF threshold of 10, corresponding to strong evidence for the presence of a QTL.

When results of both commercial populations were compared, several close significant SNPs were detected with strong effect on LG8 and LG12 in both commercial populations First, on LG8 a QTL region was identified as shared between the two commercial populations between 1,423,728 bp and 3,198,660 bp (LG8_QTL_1). This QTL explained 2.6% of the total genetic variance in pop A and 3.1% Then, several QTL regions were identified on LG12 as shared between the two populations. Based on the logBF value, 4 different QTL regions were identified: LG12_QTL_1 from 4,717,307 pb to 7,246,406 pb, LG12_QTL_2 from 8,671,217 pb to 8,797,942 pb, LG12_QTL_3 from 9,118,505 pb to 9,821,502 pb and LG12_QTL_4 from 12,167,000 pb to 19,133,722 pb. The LG12_QTL_2 QTL region was detected with a major effect in both commercial populations and this region had a length of 126,725 pb. The major SNP of this QTL region explained 21.8% of the total genetic variance in pop A and 20.3% in pop B. For all others QTL regions, the major SNPs explained a low proportion of genetic variance (less than 2%).

### LG12 QTL effect and candidate genes

Survival rates were calculated for the different genotypes of the SNP with the highest logBF value of each of LG12_QTL_2. Survival rates were studied in each of the commercial populations as well as in the validation populations. On figure 3, resistant genotypes were annotated “RR”, “RS” and “SS” for resistant, heterozygous and susceptible genotypes at LG12_QTL_2. In pop A, an average of 78% survival for the resistant genotype was reported, while this value drops to 40.6% survival with the susceptible genotype. In pop B, resistant and susceptible genotypes were associated with 66.2% and 45.4% survival, respectively. Survival rate was very similar in the validation populations, with a survival of 39.87% for the susceptible genotype while it was 63.8% for the resistant genotype. Among all other QTL regions identified on LG12, the SNP with the strongest association showed a significant impact on survival rate impact on the survival rate only in the population where it was detected, especially for QTL_LG12_1 and QTL_LG12_3 (data not shown).

**Figure 3:**
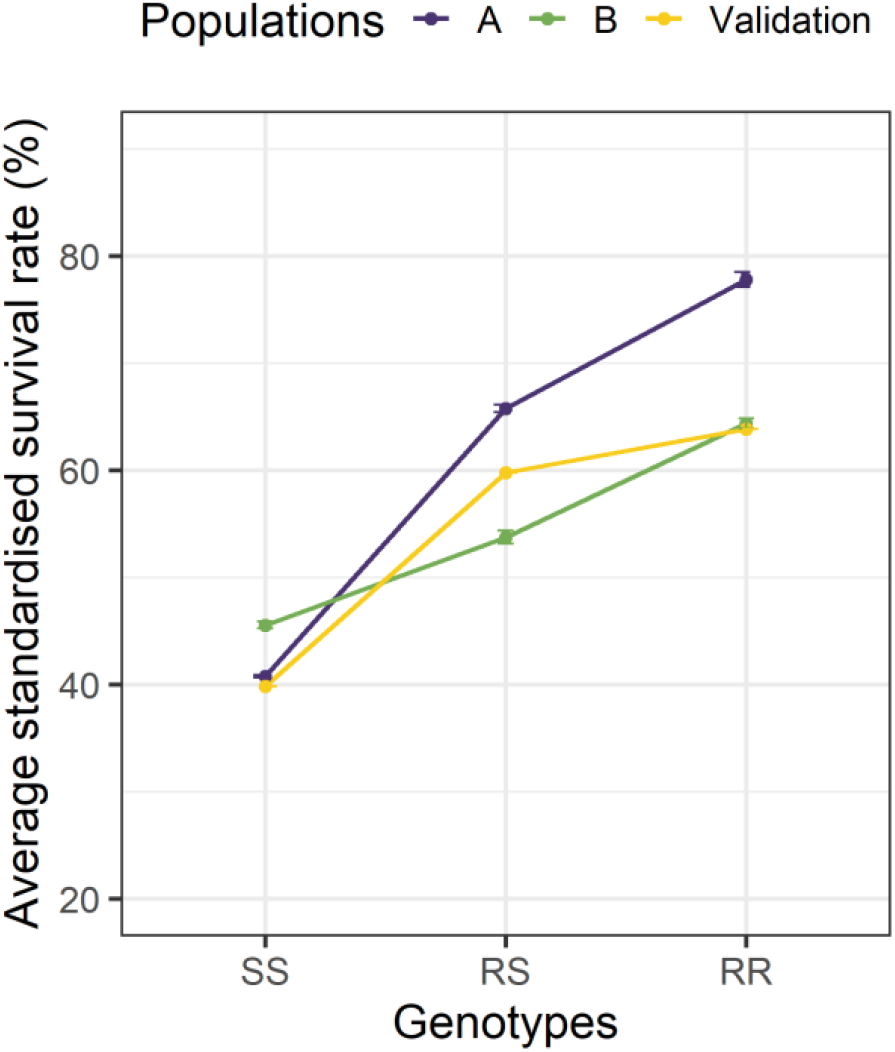
Average standardised survival rate for the SNP with higher logBF in LG12_QTL_2 detected with the BSLMM approach. Each point corresponds to a weighted mean of the average survival rate found in each commercial population cohorts. The standard deviation of survival among the three cohorts in a population was reported for each population.

The list of genes positioned in LG12_QTL_2 was obtained from the UCSC genome annotation (Figure 4). Two different genes were in that region for IFI6 coding for an interferon alpha induced protein and ZDHHC14 coding for a zinc finger palmitoyltyransferase. The IFI6 is composed of 4 exons on 903 pb and the second gene identified is ZDHHC14, characterized by 9 exons distributed on 44.4 kb. These genes are respectively positioned at 1.9 kb and 6 kb from the SNP with the highest logBF of the LG12_QTL_2.

**Figure 4:**
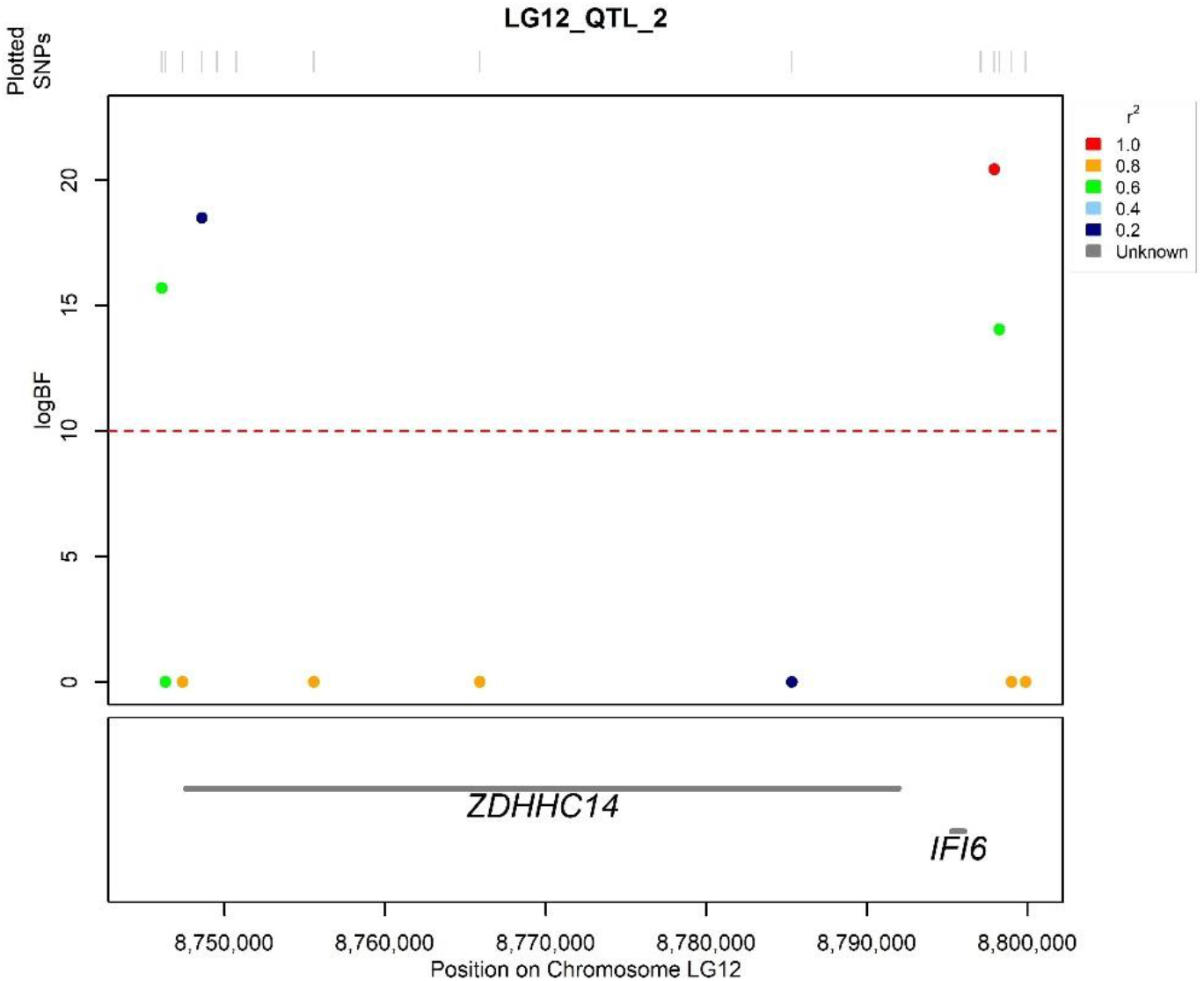
Focus on the LG12_QTL_2 region with positioning of SNPs on the genome and identification of genes in the region. The graph displays the logBF value on the y-axis and the physical position on the x-axis. Each point represents a variant of the QTL region, they are colored according to their LD r^2^ with the characteristic variant of this QTL (red point). The red dashed lines indicate the logBF=10 threshold

## Discussion

In the present study, we aimed to use a combination of whole genome sequencing, 57K genotyping, GWAS approaches to identify QTLs and then used public functional annotation to characterize these QTLs influencing VNN resistance in European sea bass.

Our results show that heritability estimates for VNN survival using imputed genotypes (0.21-0.22 on the observed scale) are of the same magnitude as values obtained in previous studies on NNV-infected sea bass, 0.08 to 0.23 on the observed scale in [10, 23] and 0.15 to 0.43 on the liability scale in [18, 39]. The number of breeders used to obtain challenged individuals as well as the number of experimental individuals, the method of infection during viral challenges, the age and size at challenge and the genetic origin and/or the experimental design are parameters that may explain the differences between the estimates of NNV survival in these studies [39].

Previous studies on the genetic architecture of VNN resistance revealed several regions characterized by the significance of one or more SNPs. The majority of the reported QTLs are putative QTLs with a defined chromosome-wide significance threshold. Through RAD sequencing, Palaiokostas et al [18] identified putative QTLs on chromosomes 3, 20 and 25 but the nomenclature used not match with the LGs used in our study. Nevertheless, after performing Blast on the European sea bass reference genome (seabass_V1.0) for the identified regions, we found that these QTLs were positioned on LG6 (for which we found no QTL) and LG12 (where we found three QTLs). Through the 57k SNP array DlabChip, Griot et al [10] identified, with a Bayesian approach, 7 putative QTL regions (LG3, LG8, LG14, LG15, LG19 and LG20) and a large QTL on LG12 in commercial populations A and B, and confirmed large QTL on LG12 in three Ifremer experimental families with an interval mapping approach. One highly significant QTL was reported to be shared between the different populations studied on LG12. More recently, Vela-Avitúa et al [23] also identified a high QTL effect in LG12 on a 7 Mb region, also using the 57k SNP array DlabChip in a different population. Thus, LG12 seems to be of special interest as a QTL was found on this chromosome in all studies and populations, except in one experimental family in [10].

Whole genome sequencing of the parents of 57K genotyped offsprings were retrieved in order to finely map SNPs and Indels to identify candidate genes for VNN resistance. Sequence variants from European sea bass genomes contain more causative mutations than the SNP genotyping arrays available for this species, making our study more robust for detecting the causal mutation. The interest of sequence data for trait mapping questions in animal breeding was uncovered in the 1000 bull genomes program [40]. In the present study, we conducted the whole-genome association studies for VNN survival using the imputed sequence data, and Bayesian fine mapping was performed to accurately map candidate variants with a binary trait. Several QTL regions were identified in the two commercial populations studied, with a majority of candidate regions revealed only in one population. Indeed, when we studied the survival rates associated to the genotypes of the majors SNPs in these regions, they had an effect on VNN survival only in the population in which they were detected. The only SNP that had a strong association with survival in both populations was the one of LG12_QTL_2. The effect of LG12_QTL_2 was further validated using the unrelated validation population where the survival rates also show a fairly significant effect linked to the resistant and susceptible genotypes at this SNP (Figure 3). The SNP characterizing this LG12 QTL shows a survival rate increasing from 40% for the susceptible genotype to nearly 80% for the resistant genotype in population A, thus marking a strong impact of this QTL in the fine mapping of the VNN resistance trait. In population B, the survival rate for the susceptible genotype is 45% but the survival rate is 64%, slightly less strong, for the resistance genotype. The figures in the validation population are 40% survival for the susceptible genotype and 64% for the resistance genotype. For the other SNPs present in the window of 127 kb, the survival rates show the same profile for the three populations as the SNP shared between the two commercial populations. In contrast, the two QTLs present at the ends of LG12 appear to be characteristic of a single population. It is therefore interesting to focus on the genomic region shared by both commercial populations. Nevertheless, finding the causal variants underlying the QTLs is a very difficult task, and therefore, very few causal variants have been identified to date. Using sequence data, we were able to refine the QTL region associated with VNN resistance and thus map the genes present at these positions. Nevertheless, prioritization of the putative causal genes is challenging with the current state of annotation of the European sea bass genome, which counts 23,382 coding genes and 35,707 transcribed genes [36].

Despite these limitations, the genes underlying the QTL region, i.e., the 127 kb region spanning the most significant SNP positions, were searched using the European sea bass genome (seabass_V1.0 [36]). The QTL region contained two genes separated by 3.6 kb, ZDNNH and IFI6 were identified as strong candidates for LG12 QTLs. The annotation of these 3 genes is conserved in several animal species and notably in fish. None of these genes have been previously reported in the literature in view of association studies for survival to VNN. Interestingly, LG12_QTL_2 and these two genes are located more than 2.5 Mb away from the QTL region identified by Vela-Avitúa et al. [23] and where the HSP70 gene is located, which was also reported in a transcriptomic analysis of response to VNN in Asian sea bass [41]. This may be a consequence of the much higher precision of our localisation, as the LG12_QTL_2 region is only 127 kb, while the region identified in [22] is 7 Mb wide. For the difference with a transcriptome study, the transcriptome identified differentially expressed genes, but in the study of [41] they compared infected and non-infected cell lines, thereby revealing genes triggered by infection, but which are not necessarily (and indeed not likely) the ones implied in disease resistance. Another difference is the way the infectious challenges are performed. Indeed, we have privileged an infection by the VNN via the contamination of the circulating water in the batch, i.e. we find the presence of the virus in the aquatic environment as encountered by the fish in rearing facilities (tanks, sea cage), whereas Vela-Avitúa et al. [23] have preferred a contamination by injection. Thus, the defence mechanisms involved in the immune response of fish may be different and therefore the gene regions highlighted by GWAS may also be different. As for the variants identified in our study, two are located in the coding region of the ZDNNH gene, which codes for a palmitoyltransferase that could catalyze the addition of palmitate onto various protein substrates. A recent study linked this reaction with the inhibition of NNV in-vitro, this study showed that protein palmitoylation and phospholipid synthesis, that involve lipid metabolism, were crucial for RGNNV replication [42]. However, the precise mechanism by which fatty acid metabolism is involved in RGNNV infection has yet to be resolved. Considering the symptoms affecting sea bass following a nodavirus infection, we can detect necrotized nerve cells with lipid droplets and also lesions in the liver and spleen tissues [15]. It is thus an interesting gene that deserves to be analysed in a more precise way via functional analyses. It is also interesting to see where the other variants of the region are positioned, and in particular the one presenting the strongest effect on the survival rate. This variant is positioned at 3,7 kb downstream of IFI6, an interferon inducible protein. This is an interesting candidate for a virus survival trait, the host responds to VNN infection through various intercellular signalling molecules, including interferon [15]. Previously, VNN infection has shown induction of interferon expression and interferon-related genes in several fish species such as Atlantic halibut, turbot, grouper, Asian sea bass and European sea bass [43–45]. In most cells, interferon response is a major first line of defense against viral infection. Viral infection triggers production of interferon, which then bind to ubiquitously expressed receptors on nearby cells and induce a powerful transcriptional program comprising hundreds of antiviral interferon stimulated genes [46]. Focusing on the IFI6 interferon inducible protein, it is involved in several processes of regulation of various viral attacks [47]. For hepatitis B virus (HBV) replication, in vivo analysis based on the hydrodynamic injection of IFI6 expression plasmid along with HBV revealed significant inhibition of HBV DNA replication and gene expression [48]. The same mechanism was also reported for the hepatitis C virus [49]. So, the resistance for VNN seems to be related to the regulation of the IFI6 protein positioned on the LG12.

Further study of the molecular mechanisms underlying the functional impact of ZNDHH disruption in-vivo is needed to better understand the role and involvement of this gene in VNN resistance. In addition, functional approaches can be conducted on the IFI6 gene to reveal expression differentials between susceptible and resistant individuals.

## Conclusions

In conclusion, we identified candidate genes associated with better resistance to VNN in European sea bass through sequence data analysis. From dense genotypic information, BSLMM approach were performed to refine association mapping analysis. Among the putative QTL identified, two QTL regions are shared between commercial populations including QTL LG12_QTL_2, where IFI6, an interferon alpha induced protein, is positioned. For practical application in selective breeding, the SNP identified in LG12_QTL_2 has a high potential to perform successful marker-assisted selection, as we showed the susceptible and the resistant genotype had a similar effect on survival to VNN infection in several, independent populations. This is a major advance to improve the resistance of cultured populations of European sea bass to one of its main challenging diseases.

## Declarations

### Consent for publication

Not applicable

## Availability of data and materials

The datasets used and/or analysed during the current study are available for scientific purposes from the corresponding author upon reasonable request.

## Authors’ contributions

ED performed the bioinformatic and statistical analyses and wrote the first draft of the paper. Aba, JB, BI and JSB organized the breeding of European sea bass. ABe and MB organized the data acquisition. ED, FA, MV, FP, ABe, MB, ABa, BI, PH and RM participated in the design of the study. MV, FP and FA provided scientific supervision. All authors read and approved the final manuscript.

## Acknowledgements

The authors would like to thank (i) the Fortior Genetics platform (ANSES, Plouzané, France) for the realization of the challenges, (ii) the INRAE genotyping platform Gentyane (INRAE, Clermont-Ferrand, France) for the production of genotype data and (iii) the GeT-PlaGe genomic platform (INRAE, Castanet-Tolosan, France).

